# The Composition of the Bacterial Community in Raw Milk from Holstein Dairy Cattle Correlated with the Occurrence of *Klebsiella pneumoniae* Clinical Mastitis Infections

**DOI:** 10.1101/2024.10.02.616256

**Authors:** Bridget O’Brien, Dongyun Jung, Soyoun Park, Daryna Kurban, Zhangbin Cai, Ngoc Sang Nguyen, Zhiwei Li, Simon Dufour, Jennifer Ronholm

**Affiliations:** Faculty of Agricultural and Environmental Sciences, Macdonald Campus, McGill University, Montréal, Québec, Canada; Mastitis Network, Saint-Hyacinthe, Québec, Canada; Faculté de médecine vétérinaire, Université de Montréal, Saint-Hyacinthe, Québec, Canada; Regroupement FRQNT Op+Lait, Saint-Hyacinthe, Québec, Canada

## Abstract

*Klebsiella pneumoniae* is a common, opportunistic bacterial pathogen that can cause severe clinical mastitis in dairy cattle. Optimizing the bovine udder microbiome to resist mastitis pathogens is a growing area of research; however, previous work has not examined which members of the mammary microbiome may have antagonistic interactions with *K. pneumoniae*. In this study, we collected quarter-level milk samples from Holstein dairy cows in Québec, Canada every two weeks for 14 months and analyzed differences in the milk microbiome between samples that were collected from healthy quarters, quarters that developed subclinical mastitis, and quarters that experienced *K. pneumoniae* clinical mastitis (KP-CM) (*n* = 512 milk samples). The occurrence of subclinical mastitis did not cause significant differences in the alpha-diversity of the milk microbiome, nor did subclinical mastitis alter the interactions between taxa in the microbiome. However, the occurrence of KP-CM caused reductions in Shannon diversity in raw milk relative to healthy milk and altered the interactions between taxa. Specifically, *K. pneumoniae* showed negative interactions with the genus *Aerococcus.* The negative interactions between *Aerococcus* spp. and *K. pneumoniae* in the context of the bovine milk microbiome should be analyzed further.

## Background

Bovine mastitis is a common disease among dairy cows that is typically caused by intramammary infections (IMIs) and results in substantial economic losses for the dairy industry [1–3]. Many possible etiological agents can cause IMI, making it a complex disease that can present in various forms. Among milk samples collected from cows with clinical mastitis (CM), 178 different species of microorganisms have been identified, but scientific literature is available for only 85 of them [4]. The etiological agents of mastitis are classified as either contagious or environmental based on their reservoirs—other cows or the environment, respectively [5].

Bovine mastitis can manifest as a subclinical or clinical form. Subclinical mastitis is more common than CM and is characterized by an elevated somatic cell count (SCC) in milk with the absence of symptoms [6,7]. The presence of symptoms defines CM which can be further categorized by a mastitis severity score (MS). Mild CM causes deviations in the milk appearance (e.g. color, consistency, viscosity) and is categorized as MS1; moderate CM is classified as MS2 and involves visible swelling of the udder in addition to abnormal milk; severe CM is classified as MS3 and includes systemic signs of infection including fever, depression, and a reduced appetite.

In contemporary dairy farming, good management practices have led to effective control of contagious mastitis pathogens, resulting in environmental mastitis becoming the most common form of mastitis on well managed farms [8]. Infections by *Escherichia coli* and *Klebsiella* spp. are common causes of environmental mastitis, and the main causes of severe CM [9,10]. Though, there are important differences between mastitis caused by *E. coli* and *Klebsiella* spp.. *E. coli* mastitis is typically short in duration and the microorganisms are usually cleared in a few days by an effective immune response, while mastitis caused by *Klebsiella* spp. is typically chronic, resulting in a longer duration of milk loss and greater risk of culling [11–13]. Among *Klebsiella* spp., *Klebsiella pneumoniae* is the most frequent species to cause CM [14]. Decreases in milk production reported after *K. pneumoniae* CM (KP-CM) may be due to damage caused to bovine mammary epithelial cells (bMEC)—the milk-producing cells in the mammary gland [15,16].

Treatment and prevention options for *K. pneumoniae* mastitis are lacking. Intramammary antibiotics are not typically recommended to treat non-severe *K. pneumoniae* infections, although, systemic antibiotic therapy is recommended for severe cases [17]. The recommendation against antibiotic treatment is not due to antimicrobial resistance (AMR), since AMR in *K. pneumoniae* isolated from bovine milk is uncommon [14,18,19], but rather the potential low efficacy of the treatment. Several studies have indicated that there may not be a benefit to treating non-severe *K. pneumoniae* mastitis with antibiotics, although further research and better treatment recommendations are needed [20,21]. Vaccines against *K. pneumoniae* mastitis are being developed and initial data look promising, but development is challenging due to the high genetic heterogeneity of *Klebsiella* isolates [22,23]. Ongoing work is also being done to develop phage cocktails to treat *K. pneumoniae* infections, although several hurdles still exist regarding this technology including the quick development of bacterial resistance and a narrow range of activity [24].

The microbiome is known to provide at least some protection against opportunistic pathogens such as *K. pneumoniae* [25], thus, optimizing the udder microbiome to resist infections has been suggested as a potential way to prevent bovine mastitis [26,27]. Numerous studies investigating the milk microbiome consistently reveal reduced diversity in milk samples from quarters with mastitis compared to healthy ones [28–30]. This reduction is often accompanied by an increased presence of reads related to the mastitis pathogen, implying an association between dysbiosis of the milk microbiome during mastitis infections. Specifically, the raw milk microbiome of *Klebsiella* spp. mastitis cases have been reported to have reduced Shannon values but no effect on the Chao1 index and a significantly lower relative abundance of Firmicutes in comparison to the milk microbiome of healthy quarters [29]. While the concept of colonization resistance has been validated for decades [25,31,32], the idea of optimizing and leveraging it to protect against bacterial pathogens deserves a closer look in the face of the AMR crisis. In this study, we hypothesize that certain bacterial commensals may be negatively correlated with the presence of *K. pneumoniae* in raw milk, and that these commensals could be used in the future for the development of anti-*K. pneumoniae* mastitis probiotics. Very few longitudinal studies have investigated the udder microbiome community with a focus on the effects of one specific mastitis pathogen. We used 16S rRNA gene amplicon and shotgun metagenomic sequencing to analyze milk samples collected throughout the lactation cycle of cows which experienced natural KP-CM and compared the results to healthy controls.

## Materials and Methods

### Milk Sample Collection

Between December 2018 and March 2020, quarter-level milk samples were collected every other week from 698 lactating Holstein cows from five commercial dairy herds in proximity to the Faculty of Veterinary Medicine of Université de Montréal (Saint-Hyacinthe, QC, Canada). These samples were collected as part of a larger study on staphylococci IMIs [4] that also examined *Staphylococcus aureus* [33] and *E. coli* mastitis [34]. In some instances, monthly sampling or opportunistic sampling of sick cattle was also performed by farmers. Farm management practices, including bedding material, dry-off protocol, feed, teat-dip product, and antibiotic treatment history are described in the supplementary material (Table S1-S3). This sampling effort led to the collection of slightly more than 27,000 milk samples. Cases of CM were identified daily by the producer or bi-weekly by the research staff and were defined by the presence of an abnormal milk sample and/or udder. Each sample was collected aseptically following the protocol developed by the Mastitis Network [35]. Briefly, a disinfectant was applied to the teat and left for the recommended contact time. Afterward, the teat was dried with a paper towel and cleaned using a disposable alcohol wipe. The cisternal milk was discarded, and approximately 50 mL of milk was then collected into a sterile Falcon tube. The samples were placed on ice and transported to the Faculty of Veterinary Medicine where they were stored at 4°C prior to being aliquoted. Aliquoted samples were frozen and stored between -10°C and -20°C until further analysis. The SCC was measured for each milk sample at Lactanet (Sainte-Anne-de-Bellevue, QC, Canada), using a flow cytometry method. Milk samples with <200,000 cells/mL were defined as healthy, while samples with >200,000 cells/mL were listed as having subclinical mastitis [36].

Microbiological culture of all milk samples was conducted following the National Mastitis Council guidelines by spreading 10uL of raw milk on 5% sheep blood agar, incubating at 35°C for 24-48h, and then enumerating different colony morphologies observed on the agar [35]. Milk samples harboring >2 dissimilar colony types on blood agar were considered contaminated according to the National Mastitis Council recommendation [35]. Colonies from non-contaminated mastitis cases were subjected to matrix-assisted laser desorption/ionization time-of-flight (MALDI-TOF) mass spectrometry to identify the etiological agent. In CM cases where a species was isolated from an infected quarter, the day that the sample was taken was considered as the day of diagnosis.

The naming convention used in this study for each cow is indicative of herd number (H), and cow traceability number (C). For example, H1C120 is a sample taken from cow 120 in herd 1.

### Sample Selection for Microbiome Study

Based on MALDI-TOF results, cows with an assigned traceability number that experienced a CM caused by only *K. pneumoniae* were selected for this study (KP-CM; *n* = 10). In CM cases where *K. pneumoniae* was isolated from an infected quarter, the day that the sample was taken for *K. pneumoniae* isolation was considered as the day of diagnosis. Milk samples (*n* = 512) from these cows were retained for DNA extraction, sequencing, and subsequent microbiome analysis. The remaining cases of KP-CM within the collection period were from cows with no traceability number recorded making it unclear which cow it occurred in or did not occur on a sampling date and therefore did not have samples from the remaining quarters collected.

### DNA Extraction

Milk samples were thawed and gently homogenized by inversion. An aliquot of 1.0mL of each milk sample was centrifuged at 16,000 x g for 10 minutes, the supernatant was discarded. DNA was extracted from pellet using DNeasy® PowerFood® Microbial Kit (QIAGEN, Germany) and the QIACube instrument (QIAGEN, Germany) following the manufacturer’s instructions.

Extracted DNA was stored at -80°C until further processing. A kit negative control was collected from each kit used in the study by extracting DNA from nuclease-free water. DNA was also extracted from a positive control, consisting of a generous donor (GD) bovine rumen sample, to assess both sample reproducibility and consistency between kits. Quality and concentration of each DNA sample was assessed using a Nanodrop 2000 (Thermo Scientific, USA) and Invitrogen™ Quant-iT™ dsDNA Assay Kit (Thermo Fisher Scientific, USA), respectively.

### 16S rRNA Gene Amplicon PCR Amplification, Library Preparation, and Sequencing

To characterize bacterial communities, the V4 hypervariable region of the 16S rRNA gene from milk samples, kit controls, and amplification negative controls, was amplified and sequenced. Amplification was performed using HotStar*Taq* Plus (QIAGEN, Germany) and the V4 index adaptor primers (F515 and R806 primer pair) in a polymerase chain reaction (PCR) [37]. The PCR included a denaturation stage (95°C, 5 minutes), 35 cycles of amplification (95℃, 30 seconds; 50℃, 30 seconds; 72℃, 1 minute), and an extension stage (72°C, 10 minutes), followed by holding samples at 4°C. Amplicons were purified using Agencourt AMPure XP (Beckman Coulter Inc., USA), then quantified using Quant-iT dsDNA Assay Kit (Invitrogen Inc., USA), as per the manufacturer’s protocol. Amplicons above 1.5ng/µL were normalized to a final concentration of 1.5ng/µL using DNA/RNA-free water, then all amplicons were pooled together. The pooled library was combined with a 15% PhiX spike-in (Illuminia Inc., USA), sequenced (251bp x 2) using the MiSeq benchtop sequencer and a 500 cycle V2 reagent kit (Illumina Inc., USA).

### 16S rRNA Gene Amplicon Sequencing Data Analysis

The FASTQ files generated from MiSeq were analyzed using Mothur (v. 1.42.3) which included taxonomic assignment using the SILVA database (v. 138) [37]. In this database, reads from the *Klebsiella* genus are listed under the taxonomic group, unclassified *Enterobacteriaceae*. Good’s coverage was calculated for each sample before rarefaction and those with coverage less than 0.99 were removed. Rarefaction and diversity metrics were done on all samples using the vegan package (v.2.6-4) in R (v.4.2.0) [38]. Samples with a library size lower than 3025 (*n* = 42) were excluded from the analysis, and the remaining samples were rarefied to a minimum library size of 3025 repeatedly (n=1000) using vegan::rarefy.perm. Samples with Good’s coverage of <0.97 after rarefaction were removed from the analysis. Beta-diversity was determined by calculating the distance between samples using vegan::vegdist with Bray-Curtis dissimilarity index, performing nonmetric multidimensional scaling (NMDS) on the calculated Bray-Curtis distances (vegan::metaMDS), then generating NMDS axis scores (vegan::scores) from the ordination and plotting calculated scores using tidyverse::ggplot2. Statistical significance of variance was determined using vegan::adonis2 with 1000 permutations. Alpha-diversity was calculated using vegan::diversity and statistical significance of alpha-diversity between groups was determined using R::wilcox.test. Differential abundance of taxa in healthy and KP-CM samples was determined using ALDEx2::aldex, which takes into account the compositional nature of the data by performing centered-log-ratio transformation on raw count data, performs Welch’s t and Wilcoxon rank test, and uses a Benjamini-Hochberg (BH) correction on raw *p*-values [39].

### Microbial Co-Occurrence Network Analysis

Co-occurrence analyses for healthy, subclinical mastitis, and KP-CM samples were performed using the Sparse Correlations for Compositional data (SparCC) method in the NetCoMi R package on raw count tables at the genera-level [40,41]. Networks were constructed using *NetCoMi::netConstruct* which utilized all genera recorded in the sample set and included very significant interactions (Correlation Coefficient ≥ +/- 0.5, *p* < 0.05). Subclinical mastitis and KP-CM networks were independently compared to healthy networks using *NetCoMi::netConstruct* with all genera and *NetCoMi::netCompare*. Differential network analysis produces Jaccard indices which assess differences between most central nodes as well as an adjusted rand index (ARI) which calculates similarity between clustering. Both a Jaccard index and ARI of 1 suggests high similarity, while an index of zero suggests that the network set has no common members.

### Shotgun Metagenomic Sequencing Preparation and Sequencing

Two DNA samples from one quarter which experienced KP-CM were selected for shotgun metagenomic sequencing. The selected samples included the sample collected at diagnosis of the KP-CM and a milk sample obtained two weeks from the same quarter before the symptoms of CM were noted. For metagenomic sequencing, DNA was extracted from 1.0 to 7.0mL of milk using DNeasy® PowerFood® Microbial Kit (QIAGEN, Germany), then cleaned using DNeasy® PowerClean® Pro Cleanup Kit (QIAGEN, Germany). Quality and concentration of each DNA sample was assessed using a Nanodrop 2000 (Thermo Scientific, USA) and Invitrogen™ Quant-iT™ dsDNA Assay Kit (Thermo Fisher Scientific, USA), respectively. Libraries for shotgun metagenomic sequencing were prepared with Nextera XT DNA Flex Library Preparation Kit (Illumina Inc., USA) and Nextera XT Index Kit (Ilumina Inc., USA), following instructions from the manufacturer. Libraries were sequenced using paired-end sequencing (151 bp x 2) on NovaSeq 6000 (Illumina Inc., USA) at Genome Quebec (Montreal, QC, Canada).

### Shotgun Metagenomic Sequencing Analysis

Raw FASTQ files generated from shotgun metagenomic sequencing were analyzed using the metaWRAP pipeline (v.1.2.1) [42]. Briefly, using the READ-QC module, reads were trimmed of adapter sequences and low-quality base pairs, and de-contaminated of host sequences using Bos taurus 3.1 bovine genome as reference (UMD 3.1, https://bovinegenome.elsiklab.missouri.edu/downloads/UMD_3.1). Taxonomy of cleaned reads was assigned using Kraken (v.2.1.2) and the minikraken v1 database, and relative abundance of species was estimated using Bracken (v.2.6.0) [43,44].

## Results

### *K. pneumoniae* Commonly Causes Severe CM

Throughout the milk collection period of all 698 cows, 341 cases of CM occurred. Among CM milk samples, 18 were considered contaminated based on the presence of >2 colony morphologies, 47 had no colony growth, 39 had a mixture of two colony morphologies; 10 of which produced at least one isolate unable to be identified, 34 produced two colony morphologies, but were determined to be the same species after MALDI-TOF, and 203 produced a single colony morphology; 12 of which were unable to be identified. From milk samples collected from quarters with CM during this study and where the growth of ≤ 2 colony morphologies was observed on blood agar after incubation, *K. pneumoniae* was isolated from 10.5% (29/276) of them. Of samples in which a single bacterial species was identified, 5.8% (13/225) had a severity score of 3, and *K. pneumoniae* was responsible for 77% (10/13) of cases with such severity score (Fig. 1). Additionally, samples from cows which experienced KP-CM had significantly higher SCC compared to samples from cows that did not experience KP-CM (p-value < 0.001, W = 6882930) (Fig. S3D).

**Fig. 1:**
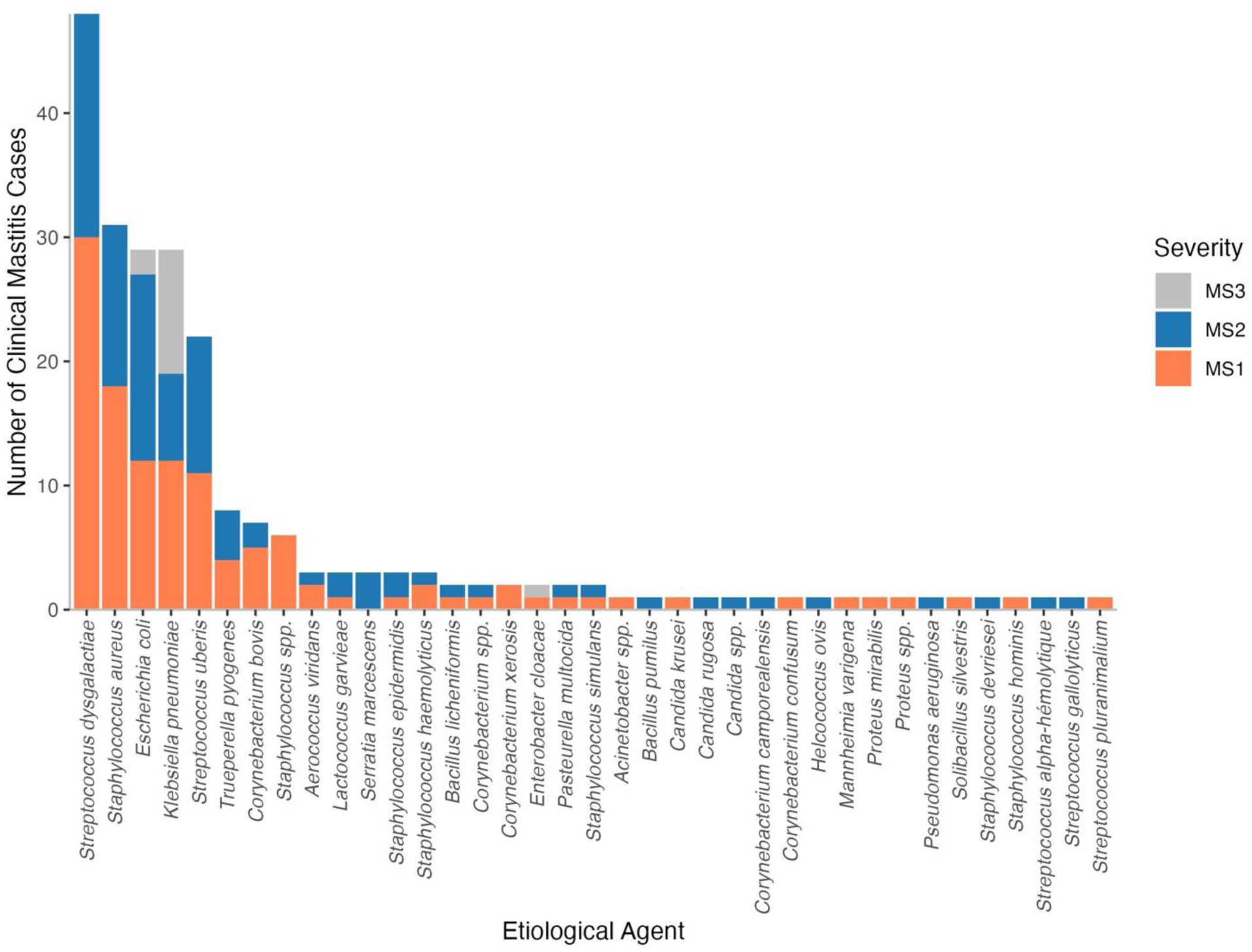
Distribution of Etiological Agents in CM Infections. The etiological agents plotted are from CM cases in which a single species was identified (*n* = 225). Of which, *K. pneumoniae* was identified in 29 of them. Proportionally, most of the severe CM cases were caused by *K. pneumoniae*.

Milk samples (*n* = 512) from 10 cows which experienced KP-CM were retained for microbiome analysis (Fig. 2). Remaining KP-CM cases were not included in our microbiome analysis because they were found through opportunistic sampling by farmers and therefore were not collected on a scheduled sampling date by the research team.

**Fig. 2.**
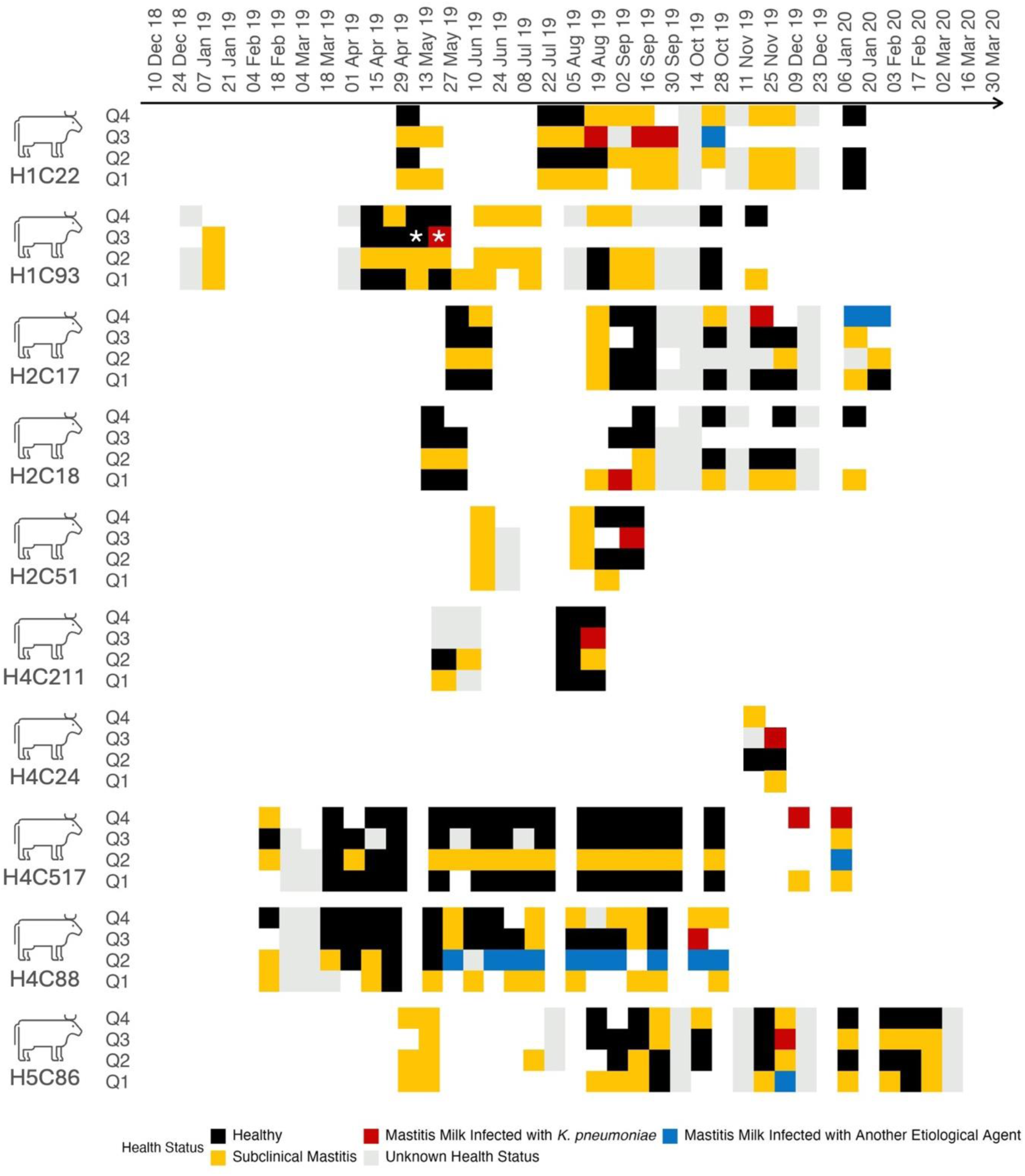
Milk Samples Collected from Holstein Cows with KP-CM. Samples collected from the 10 cows that developed KP-CM are represented in this figure. Milk samples collected from each quarter (Q1-Q4) before, during, and after the onset of KP-CM were analyzed by flow cytometry (for a SCC measurement), 16S rRNA targeted amplicon sequencing, and shotgun metagenomics. The colour of each square indicates the health status of the quarter at the time of collection. Red squares indicate a CM caused by *K. pneumoniae*, blue squares indicate CM caused by etiological agent other than *K. pneumoniae*, yellow squares indicate subclinical mastitis (SCC ≥ 200,000 cells/mL), black squares indicate healthy milk samples (SCC < 200,000 cells/mL), and grey squares indicate that the SCC level is unknown. Squares with a white asterisk represent samples which underwent shotgun metagenomic sequencing in addition to 16S rRNA amplicon sequencing.

A total of 8 milk samples could not be amplified by PCR before sequencing and were therefore removed from the study. After sequencing, a total of 24,082,364 reads passed filter, averaging to 42,324 reads per sample (samples *n* = 504, controls *n* = 65). Operational taxonomic units (OTUs) were clustered using Mothur v.1.42.3. Negative and positive controls from DNA extraction kits clustered separately (PERMANOVA: *p*=0.000999, F = 4.5704) and there was no evidence of widespread or consistent kit contamination (Fig. S1). Following quality analysis, 45 milk samples with either low (< 3025) sequence read sizes or poor Good’s coverage (< 0.99) were removed, leaving 459 milk samples for rarefaction. Milk samples (*n* = 459) were rarified using 1000 permutations. Following rarefaction, 10 milk samples had Good’s coverage < 0.97 and were therefore removed, leaving 449 milk samples for final analysis (Table S4 and Table S5). The entire microbiome analysis was conducted including and omitting milk samples considered contaminated by having ≥ 2 colony morphologies after incubation, and the same conclusions were made. Therefore, milk samples considered contaminated are included in the results presented.

Among the milk samples from cows infected with *K. pneumoniae*, 15/449 samples were from quarters with CM caused by other etiological agents, including *S. aureus*, *E. coli*, and *Lactococcus garvieae*. A total of 13/449 milk samples were taken from quarters with CM caused by *K. pneumoniae* (Table S6). Healthy samples (SCC < 200,000 cells/mL) accounted for 37% (164/449) of total samples collected, 34% (152/449) of milk samples were collected from subclinical mastitis cases (SCC ≥ 200,000 cells/mL), and 23% (105/449) of milk samples have missing SCC data making it not possible to define the health status (Fig. 2).

Taxonomic profile analysis showed that each milk sample primarily contained Firmicutes, Proteobacteria, Actinobacteriota, and Bacteroidota at different relative abundances (Fig. 3A). The relative abundance of Firmicutes varied considerably between herds, while the levels of Proteobacteria and Actinobacteriota were significantly different in herds 4 and 2, respectively (Fig. 3B). The relative abundance of Bacteroidota only differed between herds 4 and 5 (Fig. 3B). The relative abundance of unclassified *Enterobacteriaceae* was significantly higher in herd 1 than in herd 4 (Fig. S4).

**Fig. 3.**
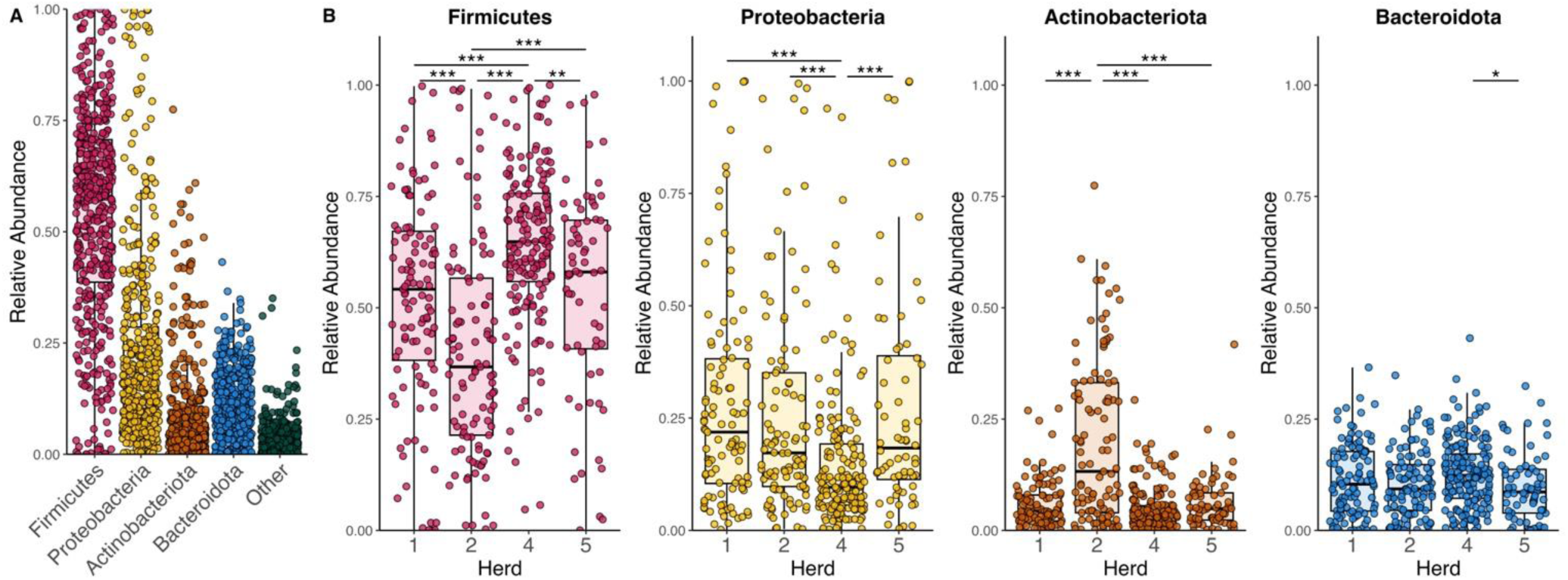
Phyla Level Variation in The Bovine Milk Microbiome. Relative abundance of bacterial taxa was identified in the raw milk microbiome of 10 cows that acquired KP-CM. (A) At the phyla level, the overall raw milk microbiome is dominated by Firmicutes and Proteobacteria. (B) The relative abundance of different phyla differs between herds, with herd 4 having a significantly lower abundance of Proteobacteria and herd 2 having a significantly higher abundance of Actinobacteria. Statistical significance was completed using a pairwise Wilcoxon test (\**p* < 0.05, \*\**p* < 0.01, and \*\*\**p* < 0.001).

### Reduced Shannon Diversity Precedes KP-CM

Compared to healthy samples (*n* = 164), milk samples taken two weeks before KP-CM was diagnosed (*n* = 7) had a significantly lower Shannon diversity (*p* = 0.00319, W = 935), yet no difference in Chao1 (*p* = 0.439, W = 661) (Fig. 4A and B). There was no difference in beta-diversity between healthy milk samples and milk samples taken two weeks before KP-CM was diagnosed (PERMANOVA: *p* = 0.1479, F = 1.2109) (Fig. 4C).

**Fig. 4.**
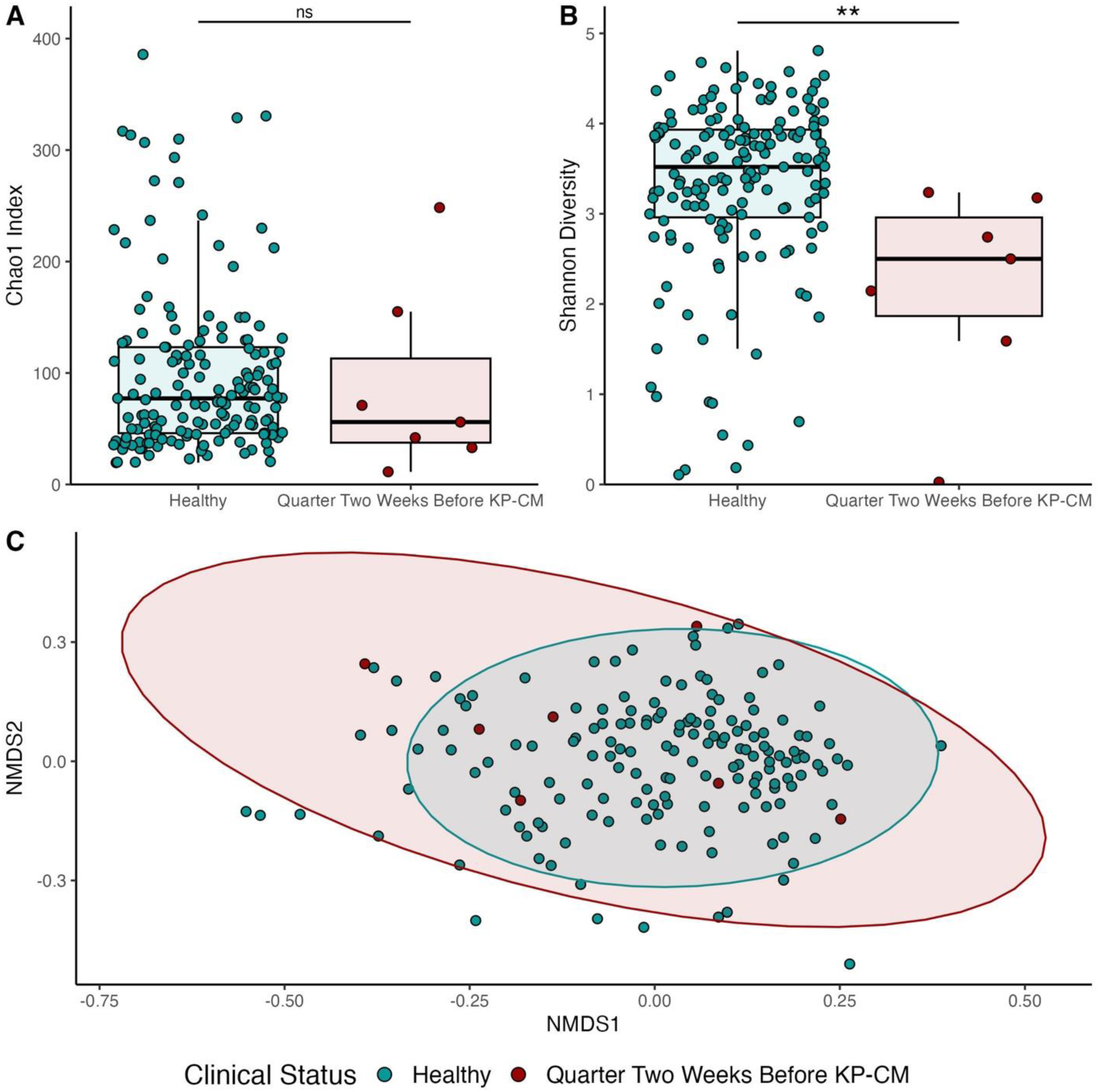
Alpha Diversity Two Weeks Before KP-CM. (A) There is no significant difference in Chao1 index between healthy samples (*n* = 164) and samples from quarters which experienced KP-CM (*n* = 7) two weeks before infection onset. (B) Significant differences were observed in Shannon diversity levels between healthy quarters and quarters which experienced KP-CM. (C) NMDS scaling analysis of Bray-Curtis distances calculated between samples showed no difference in beta-diversity two weeks before the onset of KP-CM. The statistical significance of alpha-diversity was determined by a Mann-Whitney test (\*\**p* < 0.01), and the statistical significance of beta-diversity was determined using PERMANOVA (*R::adonis2*).

### High Levels of Unclassified *Enterobacteriaceae* are Found in Milk from KP-CM quarters

At the time that KP-CM was confirmed, 7/10 cows had clinical or subclinical mastitis in at least one other quarter (Fig. 2). Samples of KP-CM in the remaining three cows (H2C46607, H2C7454, H2C27299) lacked SCC data. The beta-diversity was significantly different between healthy and KP-CM samples (PERMANOVA: *p* < 0.001, *F* = 8.0868) (Fig. 4A). Alpha-diversity metrics were also compared between healthy samples and KP-CM samples. No significant difference was observed in Chao1 index between healthy and KP-CM samples (*p* = 0.866, W = 1096), while Shannon diversity was significantly lower among KP-CM samples (*p* = 0.003647, W = 1583) (Fig. 5B).

**Fig. 5.**
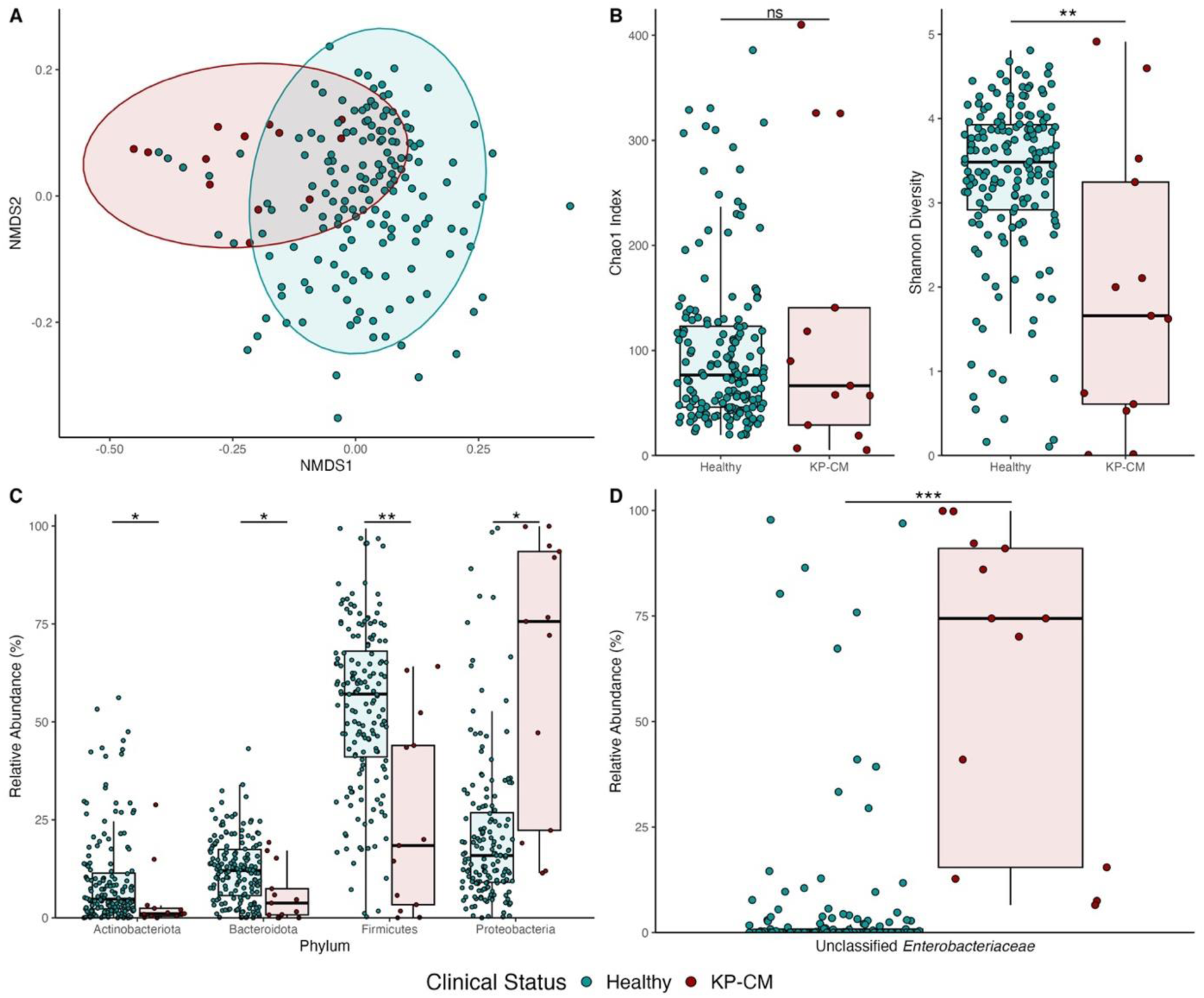
Diversity Metrics and Relative Taxonomic Abundance in Healthy and KP-CM Samples. (A) NMDS scaling analysis of Bray-Curtis distances calculated between healthy (*n* = 164) and KP-CM (*n* = 13) samples. (B) Alpha diversity metrics between healthy and KP-CM samples reveal no significant difference in Chao1 index between groups, while Shannon index is significantly reduced in KP-CM samples compared to healthy. Statistical significance was determined by a Mann-Whitney test (\*\**p* < 0.01). (C) Significant increases in the relative abundance of Proteobacteria, and significant decreases in Firmicutes, Bacteroidetes, and Actinobacteriota are observed in KP-CM samples, compared to healthy milk samples. (D) Among genera, only significant increases in unclassified *Enterobacteriaceae* can be seen in KP-CM samples. Differential abundance analysis at both phylum and genera level was completed using ALDEx2 with a BH corrected p-value (\**p* < 0.05, \*\**p* < 0.01, and \*\*\**p* < 0.001).

Differences between healthy and KP-CM samples were observed by the variance in relative abundance of taxa. Differential abundance was carried out using ALDEx2 which considers the compositional nature of high-throughput sequencing data. Results showed significant reductions in Firmicutes (*p* = 0.00374), Actinobacteriota (*p* = 0.0203), and Bacteroidota (*p* = 0.0248), as well as significant increases in Proteobacteria (*p* = 0.00192) in KP-CM samples (Fig. 5B). Among Proteobacteria, significant increases in unclassified *Enterobacteriaceae* among KP-CM cases in comparison to healthy quarters were observed (Fig. 5C) (*p* < 0.0001). This suggests that *K. pneumoniae* dominates other members of the microbiome during infection.

### Alpha-Diversity is Unaffected by Subclinical Mastitis

Since this study had a high number of samples with subclinical mastitis, we wanted to further investigate the differences between healthy (*n* = 164) and subclinical mastitis (*n* = 152) samples to determine if subclinical mastitis affects the diversity of the microbiome in a similar way to KP-CM. Analysis of alpha diversity metrics revealed no differences in Chao1 (*p* = 0.4467, W = 11846) or Shannon Diversity (*p* = 0.07557, W = 13906) (Fig. 6A). No obvious clustering can be observed between groups when plotting NMDS1 and NMDS2 of healthy samples and subclinical mastitis samples, though statistical significance was recorded (PERMANOVA: *p* < 0.001, *F* = 4.1465) (Fig. 6B). This indicates that the overall composition of the microbiome is not affected in subclinical mastitis as it is in CM.

**Fig. 6.**
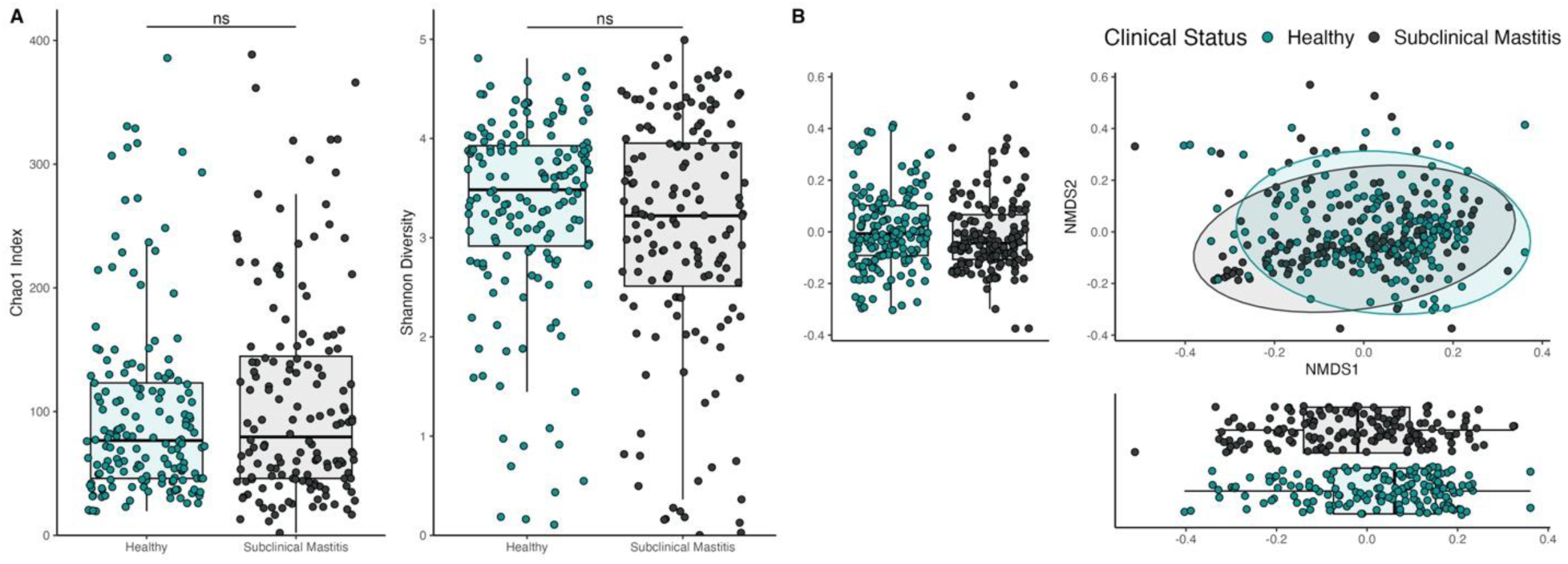
Diversity Metrics in Subclinical Mastitis and Healthy Samples. (A) Alpha diversity metrics between healthy and subclinical mastitis samples reveal no significant difference in Chao1 index or Shannon diversity between healthy and subclinical mastitis. Statistical significance was determined by a Mann-Whitney test. (B) NMDS plot of healthy (*n* = 164) and subclinical mastitis (*n* = 152) samples. Beta-diversity was dissimilar between healthy and subclinical mastitis samples (PERMANOVA: *p* < 0.001, *F* = 4.1465).

### KP-CM Changes Interactions Between Taxa in the Milk Microbiome

Co-occurrence networks using KP-CM (*n* = 13), subclinical mastitis (n =152), and healthy samples (*n* = 164) were generated to understand interactions between bacteria in different clinical settings. The adjusted Rand index (ARI) which looks at the similarity between clustering, indicates high similarity between networks composed of healthy and subclinical mastitis samples (ARI = 0.773, p-value = 0.00), while a lower degree of similarity between healthy and KP-CM networks (ARI = 0.131, p-value = 0.00) (Fig. 7). Furthermore, Jaccard indices, which looks at the similarity between sets of the most central nodes and hub nodes between the two centers, highlighted several significant differences between healthy and KP-CM networks, while no significant differences between healthy and subclinical mastitis networks (Table 2). This suggests that KP-CM alters interactions between taxa within a healthy milk microbiome, while subclinical mastitis does not.

**Fig. 7.**
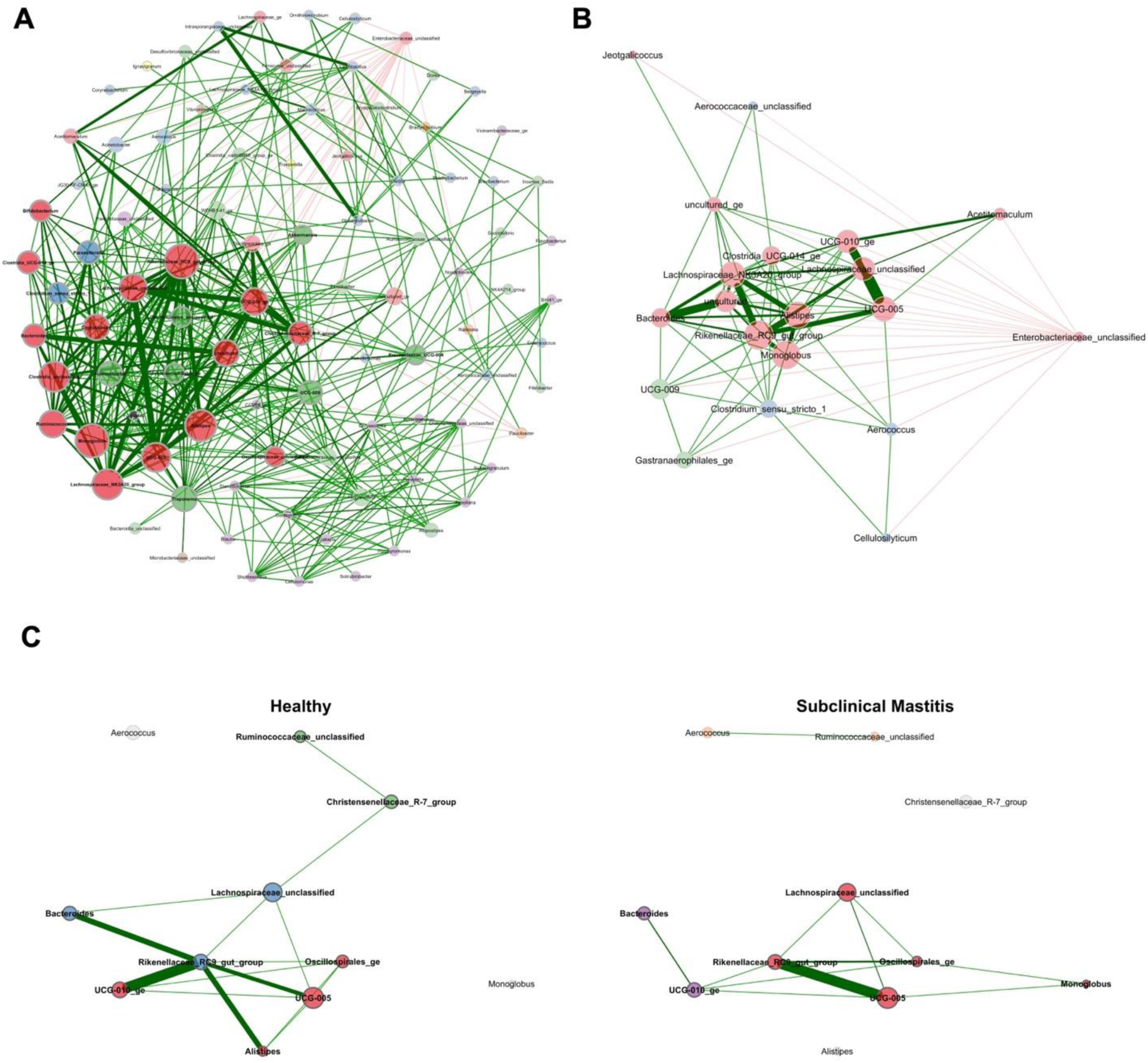
Microbial Co-Occurrence Networks of KP-CM, Healthy, and Subclinical Mastitis Samples. Microbial co-occurrence networks were constructed using all genera recorded genera in the 16S rRNA gene sequencing sample set. Each node represents one genus and the color of the edge which connects nodes denotes positive (green; *r* > 0.5) or negative (red; *r* < -0.5) interactions, with the thickness of the edge representing the strength of their association. The size of each node is proportional to its eigenvector centrality (a measure of influence a node has in a network), and its color represents its cluster. (A) Microbial co-occurrence network of KP-CM (*n* = 13) samples is composed of eight clusters. (B) Unclassified *Enterobacteriaceae* is recorded to have only strong negative interactions with other genera of the milk microbiome. (C) Healthy (*n* = 164) and subclinical mastitis (*n* = 152) networks both have few significant interactions between genera.

**Table 2:** Jaccard index values for microbial co-occurrence networks.

| Comparison Between Healthy and KP-CM Networks |  |  |  |
| --- | --- | --- | --- |
| Parameters | Jacc | P (≤ Jacc) | P (≥ Jacc) |
| Degree | 0.082 | 0.000000 *** | 1.000000 . |
| Betweenness centr. | 0.020 | 0.000000 *** | 1.000000 . |
| Closeness centr. | 0.082 | 0.000000 *** | 1.000000 . |
| Eigenvector centr. | 0.082 | 0.000000 *** | 1.000000 . |
| Hub taxa | 0.273 | 0.361971 . | 0.793851 . |
| Comparison Between Healthy and Subclinical Mastitis Networks |  |  |  |
| Parameters | Jacc | P (≤ Jacc) | P (≥ Jacc) |
| Degree | 0.600 | 0.980338 . | 0.076564 . |
| Betweenness centr. | 0.167 | 0.351166 . | 0.912209 . |
| Closeness centr. | 0.600 | 0.980338 . | 0.076564 . |
| Eigenvector centr. | 0.600 | 0.980338 . | 0.076564 . |
| Hub taxa | 0.600 | 0.980338 . | 0.076564 . |

Significance codes: \*\*\*: $P \leq 0.001$ , \*\*: $P \leq 0.01$ , \*: $P \leq 0.05$ , .: $P \leq 0.1$

In the KP-CM network, a pattern of interactions distinct from healthy and subclinical mastitis networks can be observed. Unclassified *Enterobacteriaceae* was shown to have only negative interactions with other taxa, including *Aerococcus* (*r* = -0.51) and unclassified *Aerococcaceae* (*r* = -0.67). In addition, unclassified *Enterobacteriaceae* had strong negative interactions with most genera (6/9) reported in the network composed of healthy samples (Alistipes; *r* = -0.67, UCG-005; -0.58, UCG-010_ge; -0.62, Rikenellaceae_RC9_gut_group; *r* = - 0.58, Lachnospiraceae_unclassified; r = -0.61; Bacteroides; r = -0.63; (Fig. 7A and B). This suggests that *K. pneumoniae* may interact negatively with the core taxa of a microbiome of milk from a healthy quarter to establish infections, and *Aerococcus* may interact negatively with *K. pneumoniae* to prevent the occurrence of KP-CM.

### Aerococcus urinaeequi or Aerococcus viridans may prevent the onset of KP-CM

To further understand interactions occurring between taxa before and during KP-CM, shotgun metagenomic sequencing was performed on a healthy milk sample collected two weeks before the onset of KP-CM as well as during KP-CM from H1C93. Among samples, 122,336,818 reads passed filter, averaging to 61,168,409 reads per sample. Following the removal of host DNA, 1,521,294 reads remained amongst the samples, averaging to 760,647 reads per sample.

In accordance with the network analysis, *Aerococcus,* a genus which unclassified *Enterobacteriaceae* had strong negative interactions with, was identified in a healthy milk sample two weeks before the onset of KP-CM. Within this sample, 24% of bacterial reads were assigned *Aerococcus*. *Aerococcus urinaeequi* composed ∼21% of bacterial reads, while ∼3% of bacterial reads were classified as *Aerococcus viridans*. In the same sample, *K. pneumoniae* was mapped to ∼5% of reads. Due to the immense amount of host DNA sequenced in samples, little reads could be mapped to functional genes and no metagenome-assembled genomes in the healthy milk sample could be generated, making the potential mechanism of antagonism between *Aerococcus* spp. and *K. pneumoniae* unclear. Alongside the negative interactions noted to occur between *Aerococcus* spp. and *K. pneumoniae* in the KP-CM co-occurrence network, this result reveals that *K. pneumoniae* may be interacting negatively with specifically *Aerococcus urinaeequi* or *Aerococcus viridans*.

Within the sample from the KP-CM case, the majority (97%) of bacterial reads were mapped to *K. pneumoniae* and no reads were mapped to *Aerococcus*. This result further suggests that *K. pneumoniae* displaces *Aeroccocus* and other taxa during to the onset of CM (Fig. 8).

**Fig. 8.**
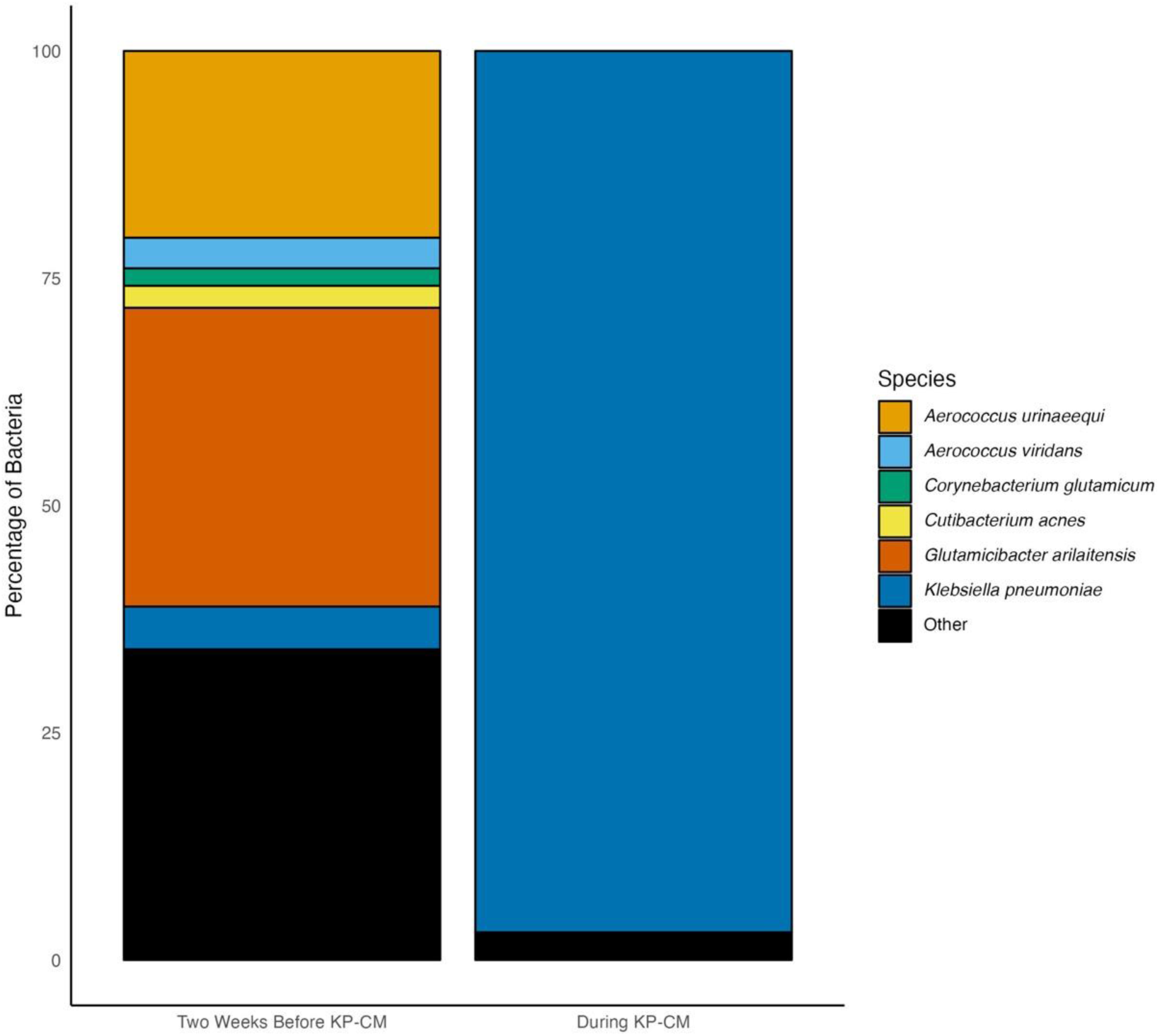
Species Relative Abundance Before and During KP-CM in H1C93. Shotgun metagenomic sequencing identified the presence of *A. urinaeequi*, *A. viridans*, and *K. pneumoniae* two weeks before KP-CM in H1C93. During KP-CM, *K. pneumoniae* is the main species identified with no *Aerococcus* species present.

## Discussion

Mastitis can be caused by several bacterial pathogens, each of which have unique virulence factors that interact with the core microbiota differently. Understanding the microbiome composition and health status of cows which experienced KP-CM can aid in understanding what predisposes cows to this infection and ultimately how to prevent them from occurring. In this study, we attempted to characterize the milk microbiome of cows infected with *K. pneumoniae* by performing 16S rRNA gene amplicon sequencing and shotgun metagenomic sequencing on milk samples collected before and during a natural KP-CM. Our results indicate that *K. pneumoniae* is an opportunistic pathogen that tends to infect cows with high SCC and quarters with reduced diversity. During KP-CM, *K. pneumoniae* interacts negatively to displace resident taxa of raw milk, and we predict *Aerococcus* spp., specifically *A. urinaeequi* or *A. viridans,* may have antagonistic activity against *K. pneumoniae*.

Cows within this study, had high inflammation throughout lactation cycles (Fig. S3), with most milk samples of known SCC indicating the occurrence of subclinical mastitis and, therefore, stress on the udder [45]. Furthermore, two weeks before the onset of KP-CM, milk samples from quarters soon to be infected with *K. pneumoniae* had reduced Shannon diversity, suggesting that in addition to cows that commonly experience subclinical mastitis, quarters with low diversity may be more susceptible to *K. pneumoniae* mastitis. In humans, classical *K. pneumoniae* is an opportunistic pathogen capable of causing infections in immunocompromised or unhealthy individuals [46]. Opportunistic pathogens are often thought to be commensal. Indeed, *K. pneumoniae* is frequently detected in the rumen, bedding, water, and manure on dairy farms [47,48]. Given that *Klebsiella* spp. are recognized as environmental pathogens, *K. pneumoniae* may not be commensal of the udder. In most healthy samples (73%; 124/164), unclassified *Enterobacteriaceae* was in very low relative abundance (< 1%), and it had no interactions in the network analysis of samples from healthy quarters, suggesting that it is not frequently found in high abundance within a healthy milk microbiome. Some healthy milk samples contained high levels of unclassified *Enterobacteriaceae* and may have been a result of environmental contamination, insufficient sequencing depth, or represented the beginning of an infection with a delay in the host immune response.

Compared to healthy samples, samples from KP-CM cases had reduced Shannon values, but no effect on the Chao1 index, and an increased relative abundance of Proteobacteria, which is in agreement with a previous study [29]. This suggests that the richness of the milk microbiome seems to be unaffected by KP-CM, but the evenness is. Indeed, an ALDEx2 analysis, Wilcoxon rank-sum (data not shown), and LEfSe analysis (data not shown) indicated there were significant increases in unclassified *Enterobacteriaceae* among KP-CM cases compared to healthy. Other studies have shown that *K. pneumoniae* appears to dominate the microbiome in other infections, including in the gut microbiota of human patients with bacteremia [49] and sepsis [50].

It is not known how *K. pneumoniae* dominates microbiomes. Research on interbacterial interactions with *K. pneumoniae* is sparse, making it unclear if a distinct interaction pattern may occur during KP-CM. In hospital settings, acquisition of *K. pneumoniae* infections is commonly associated with cross-contamination in *K. pneumoniae*-prevalent environments [51]. Moreover, during *K. pneumoniae* infections in immunocompromised patients, intestinal colonization with *K. pneumoniae* was found to be significantly associated with subsequent infections [52,53]. A similar case can be made for dairy farming. It is known that *K. pneumoniae* is shed in the feces of dairy cows and can remain in bedding material [47,54]. As a result, bedding can become a potential source of *K. pneumoniae* if not properly maintained and replenished [54].

Similar to previous studies, *Aerococcus* may represent one taxa with protective effects in the udder [33,55]. Our study was able to identify *A. urinaeequi* or *A. viridans* as the possible species, though no mechanism of action was able to be identified due to the immense amount of host DNA present in sequencing reads. However, one study has shown that a purified antibacterial substance from *A. urinaeequi* is inhibitory towards *K. pneumoniae* growth *in vitro* [56]. The relationship between *Aerococcus* and mastitis-causing species, including *K. pneumoniae*, should be further investigated to understand if interbacterial interactions occurring in the udder can be leveraged to prevent infection.

There was no difference in alpha diversity or network analyses in milk samples from healthy quarters and quarters with subclinical mastitis. The lack of difference in alpha diversity or microbial network interactions between healthy and subclinical mastitis samples suggests that dysbiosis may be associated with the onset of symptoms. Perhaps the increase in SCC is a result of an effective immune response, capable of keeping levels of the pathogen low, thus preventing such dysbiosis from occurring and symptoms presenting.

In this study, we investigated the bovine udder microbiome of Holstein cows with KP-CM infections. Although limited by small sample size, our results show that similar to in human infections, *K. pneumoniae* acts as an opportunistic pathogen among dairy cows, causing CM in cows which frequently experience subclinical mastitis and in quarters with low alpha diversity. *K. pneumoniae* is a commensal in the dairy farm environment, and therefore infection may result from cross-contamination with the cows’ surrounding materials. During infection, *K. pneumoniae* dominates the microbiota overgrowing other bacteria such as *Aerococcus*, resulting in reduced alpha diversity in infected quarters. However, subclinical mastitis cases did not have reduced alpha diversity in comparison to samples from healthy quarters, and thus symptom onset may be associated with microbial dysbiosis. Overall, reducing the prevalence of *K. pneumoniae* within the environment of the dairy farm may be an essential strategy for preventing the chance of cross-contamination and, consequently, the occurrences of KP-CM. Further studies are necessary to understand the relationship between *Aerococcus sp*. and *K. pneumoniae*, and if *Aerococcus sp*. can be leveraged to prevent infection.

## Data Availability

All sequencing data from milk samples were deposited in the Sequence Read Archive (SRA) under Bioproject PRJNA948368.

## Author Contributions

JR and SD: conceived of the study, experimental design and funding acquisition. SD and DK collected milk samples, plated samples on blood agar, and identified the causative agents of mastitis. BO: extracted DNA from milk samples, conducted bioinformatic analysis of 16S rRNA sequences, performed shotgun metagenomic sequencing, and analyzed the data. BO, DJ, SP, ZC, SN and ZL performed sequencing of 16S rRNA DNA. BO: wrote original draft; all contributing authors: writing, reviewing, and editing.

## Acknowledgements

This research was undertaken, in part, thanks to funding from the Canada Research Chairs Program.

